# Decision-making increases episodic memory via post-encoding consolidation

**DOI:** 10.1101/311571

**Authors:** Vishnu P. Murty, Sarah DuBrow, Lila Davachi

## Abstract

The ability for individuals to actively make decisions engages regions within the mesolimbic system and enhances memory for chosen items. In other behavioral contexts, mesolimbic engagement has been shown to enhance episodic memory by supporting consolidation. However, research has yet to investigate how consolidation may support interactions between decision-making and episodic memory. Across two studies, participants encoded items that were occluded by cover screens and could either actively decide which of two items to uncover or were pre-selected by the experimenter. In Study 1, we show that active decision-making reduces forgetting rates across an immediate and 24-hour memory test, a behavioral marker of consolidation. In Study 2, we use functional neuroimaging to characterize putative neural markers of memory consolidation by measuring post-encoding interactions between the hippocampus and perirhinal cortex (PRC). We show that choice-related striatal engagement is associated with increased post-encoding hippocampal-PRC interactions. Finally, we show that a previous reported relationship between choice-related striatal engagement and long-term memory is accounted for by post-encoding hippocampal-PRC interactions. Together these findings support a model by which actively deciding to encode information enhances subsequent consolidation mechanisms to preserve episodic memory for outcomes.

## Introduction

Individuals value the ability to actively make decisions and manipulate their environment (Leotti, Iyengar, & Ochsner, 2010). While previous research has characterized the prioritization of valuable information in episodic memory (Miendlarzewska, Bavelier, & Schwartz, 2016; Murty & Adcock, 2017), these processes have not been fully characterized in the context of decision-making. Recent research has shown that the simple act of making a decision enhances episodic memory by engaging hippocampal and mesolimbic systems during choice and encoding (Murty, DuBrow, & Davachi, 2015). In parallel, animal research has shown that engagement of mesolimbic systems during encoding supports episodic memory by strengthening post-encoding consolidation (Wang & Morris, 2010). However, research has yet to fully characterize how post-encoding memory consolidation processes may be initiated by active decision-making. In the current study, we used behavioral and neural markers of postencoding consolidation to characterize a novel mechanism by which decision-making enhances episodic memory.

The opportunity to actively make decisions and implement agency over one’s environment engages regions associated with mesolimbic dopamine systems. When participants are given the opportunity to choose which of two gambles to partake in there is increased engagement of the striatum and dopaminergic midbrain (Leotti & Delgado, 2011, 2014). We recently showed a parallel mechanism is engaged when individuals are given the opportunity to make decisions about what information to encode (Murty et al., 2015). In this study, we manipulated whether participants could actively decide which information to learn. Memory was enhanced for items selected by the participants versus items selected by the experimenter, and these memory enhancements were related to striatal engagement during choice behavior. Together these studies suggest that the act of decision making enhances striatal activation—a proxy of mesolimbic engagement—and episodic memory.

Rodent and human studies have shown that mesolimbic engagement enhances memory, in part, by increasing memory consolidation. Memory consolidation in this context refers to the strengthening of memory after encoding and prior to retrieval, often demonstrated as a resistance to forgetting over time. Behavioral studies have shown that memory enhancements of information encoded under reward motivation, which is thought to engage mesolimbic dopamine systems, only emerge after a significant delay (Murayama & Kitagami, 2014; Murayama & Kuhbandner, 2011; Patil, Murty, Dunsmoor, Phelps, & Davachi, 2017). Further, rodent studies show that reward and novelty-based enhancements rely on dopamine-mediated consolidation (Abraham, Neve, & Lattal, 2016; Li, Cullen, Anwyl, & Rowan, 2003a; Salvetti, Morris, & Wang, 2014; Takeuchi et al., 2016; Wang & Morris, 2010; Wang, Redondo, & Morris, 2010). Thus, if the same mechanism that enhances memory in these contexts also engages the mesolimbic system during decision-making, then previously identified decision-induced memory enhancements may upregulate post-choice consolidation processes.

Behavioral and neuroimaging research have identified multiple methodological approaches to characterize memory consolidation in humans. Behaviorally, memory consolidation can be measured by testing memory both at immediate and delayed time points. Importantly, querying memory at both time points allows one to compute the slope of forgetting, which should be reduced for memories strengthened by postencoding consolidation processes. Neurally, memory consolidation can be characterized by relating episodic memory with measures of offline, post-encoding activity. Systems memory consolidation is thought to transfer information initially encoded in the hippocampus to cortical regions (McClelland, McNaughton, & O’Reilly, 1995; Nadel, Samsonovich, Ryan, & Moscovitch, 2000). Thus, one signature of this process would be increased memory network coupling following encoding. In line with this framework, neuroimaging studies have shown that increased functional coupling between hippocampus and category-selective regions predicts delayed memory performance (Murty, Tompary, Adcock, & Davachi, 2017; Schlichting & Preston, 2016; Tambini, Ketz, & Davachi, 2010; Tompary, Duncan, & Davachi, 2015). In the current study, we characterized these behavioral and neural markers of post-encoding consolidation and their relationship to decision-related memory enhancements.

Across two studies, participants completed a choice memory paradigm in which we manipulated whether participants could either actively decide which information to encode (Choice) or the information was selected for them (Fixed, Figure 1). The choice condition imbued participants with a sense of agency over their environment, but critically, had no effect on which memoranda were shown. In Study 1, we characterized behavioral markers of consolidation by testing forgetting rates for items actively selected by the researcher (Choice) versus items selected by the experimenter (Fixed). In Study 2, we characterized neural markers of consolidation by measuring post-encoding changes in functional coupling between the hippocampus and the perirhinal cortex (PRC). We selected the PRC as our cortical target given its critical role in mediating memory for object images (Awipi & Davachi, 2008; Davachi, 2006; Davachi, Mitchell, & Wagner, 2003; Graham, Barense, & Lee, 2010; L. Litman, Awipi, & Davachi, 2009; Staresina, Duncan, & Davachi, 2011), and prior work implicating hippocampal-PRC coupling as a neural marker for object based memory consolidation (Vilberg & Davachi, 2013). Specifically, we tested whether hippocampal-PRC connectivity was (1) enhanced after encoding, (2) related to engagement of the striatum during decision-making, and (3) related to 24-hour memory for actively selected (Choice) items.

**Figure 1:**
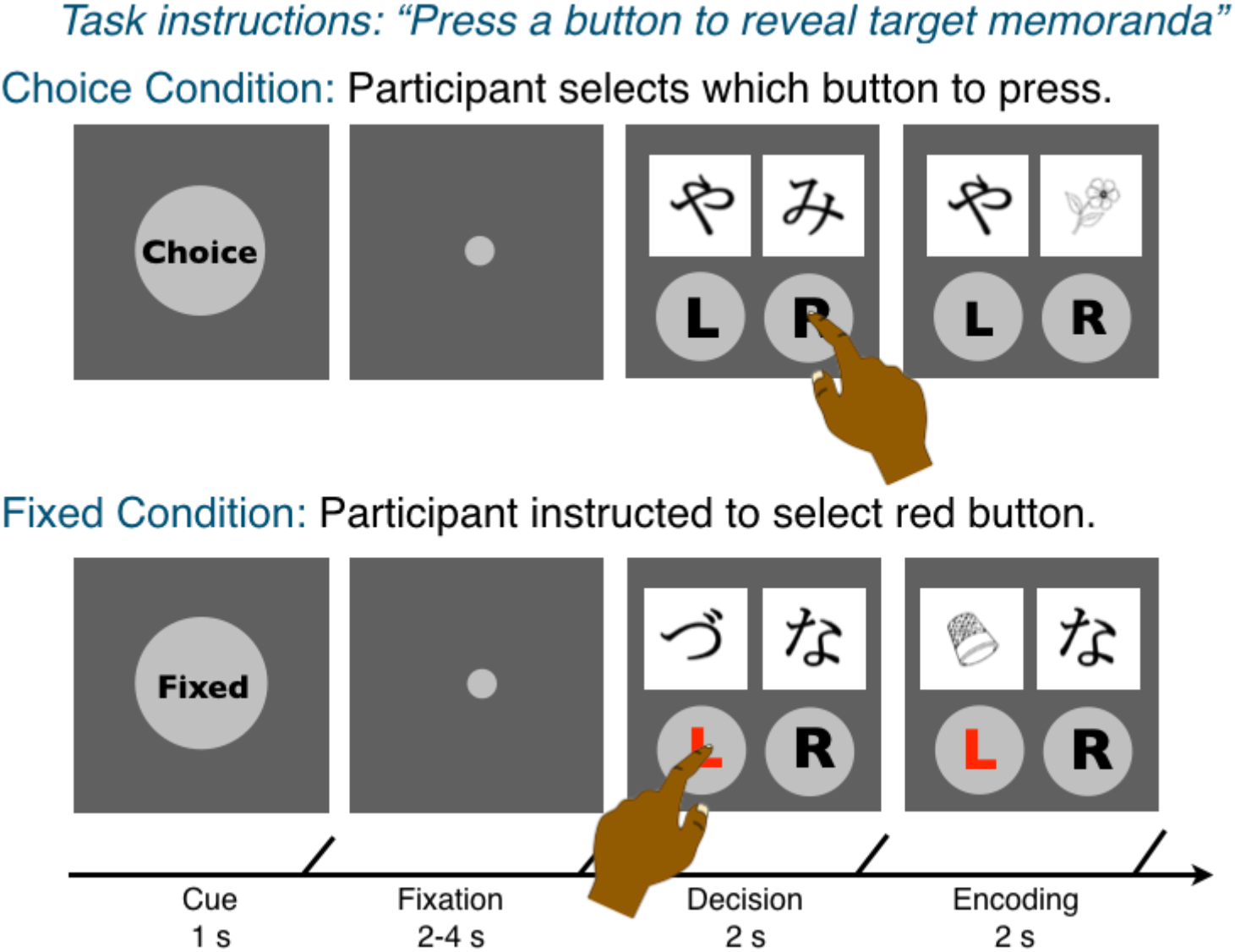
Choice Memory Encoding Task

## Methods

### Participants

Participants were recruited from the New York University and New York City communities. Informed consent was obtained for each participant in a manner approved by the University Committee on Activities Involving Human Subjects. In Study 1 (behavioral), 36 healthy, right-handed participants were paid $25 to participate. Three participants were excluded due to failure to follow task instructions (n=1), familiarity with the stimuli (n=1), and failure to complete the 24-hour memory recognition test (n=1). The final sample included 33 participants (21 females; 18-35; median age = 22 years). In Study 2 (fMRI), 24 healthy, right-handed participants were paid $50 to participate. 4 participants were excluded due to failure to follow task instructions (n=1), poor neuroimaging data quality (n=1), and failure to complete the 24-hour memory recognition test (n=2). Portions of this data relating to memory encoding, but not consolidation, have been reported elsewhere (Murty et al., 2015).

### Behavioral Paradigm

Participants in both studies completed a multi-phase choice memory task, which probes how arbitrary decision making—giving individuals the opportunity to make a choice—influences episodic memory. The task consists of 4 phases: 1) pre-encoding ratings, 2) choice memory encoding, 3) post-encoding ratings, and 4) a memory test. In the pre-encoding ratings phase, individuals made preference ratings about 80 hiragana characters. They were instructed to indicate how much they ‘liked’ each character on a 5-point scale. Participants had 4 seconds to rate each character (2-5 s ITI). 60 of the most neutrally related characters from the pre-encoding ratings phase were used as cover screens for the choice encoding task (see Murty et al., 2015 for more details). Next, participants completed the choice phase (Figure 1), which manipulated whether individuals could actively make decisions about which information to encode. On each trial, participants were first shown a cue for 1 s indicating the condition (i.e., choice, fixed), followed by a fixation dot for 2-4 seconds, followed by a decision phase for 2 seconds, and an encoding phase for 2 seconds.

During the decision phase, participants saw a screen with two cover screens, which were previously rated hiragana characters, and two buttons underneath. Participants were instructed to make a button press to reveal an object occluded by the cover screen. During encoding phase, participants were instructed to encode the previously occluded object image. In the choice condition, participants actively choose which cover screen to remove. In the fixed condition, the participants were instructed to choose the button that was highlighted with red text. If participants took longer than 2 seconds to respond, they were shown a screen indicating their response was too slow and no image was presented. Unbeknownst the participants, object images were pre-selected, so there was no relationship between decisions and the underlying object image. Following each trial, a fixation cross appeared for 3-24 seconds in a manner optimized for fMRI analysis (https://surfer.nmr.mgh.harvard.edu/optseq/). Participants completed 60 choice trials and 60 fixed trials intermixed across four 8-minute runs. Following this choice memory encoding phase, individuals completed the post-encoding ratings. This session was identical to the pre-encoding ratings task. Finally, participants completed the memory test phase, a self-paced recognition test for the object images presented during the encoding task. Participants were shown object items one at a time and had to indicate whether they previously viewed the object (yes/no), and the confidence in their response (“very sure”, “pretty sure”, “just guessing”). Trial duration was self-paced (1s ITI). Participants completed 240 recognition memory trials including 60 objects from the choice condition, 60 objects from the fixed condition, and 120 novel/foil objects. In study 1, recognition memory was split over two sessions (immediate memory test, 24-hour memory test; detailed below), with half of the items in each condition (choice, fixed and novel) appearing in each. In study 2, the complete recognition memory test occurred after a 24-hour delay.

### Experimental Protocol

In Study 1, participants were consented and given instructions about the study. Participants then completed the pre-ratings task, the choice memory encoding task, and post-ratings task. Participants then completed the memory test for half of the stimuli presented during encoding. Participants returned approximately 24-hours later to complete a memory test for the other half of the stimuli presented during encoding. Participants were then paid and de-briefed about the experiment. All sessions took place in a behavioral testing room. In Study 2, participants were consented and given instructions outside of the scanner. Once inside the scanner, participants completed the pre-encoding ratings, the choice memory encoding, and the postencoding ratings phase. Participants returned approximately 24-hours later to a behavioral testing room and performed the memory test for all stimuli. Participants were then paid and de-briefed about the experiment.

### Study 1: Analysis

To test whether individual’s memory was above chance, we compared the percentage of objects endorsed as old for old versus new items within-subjects for each testing day. Next, to determine if there were choice-related memory enhancements, we compared the percentage of objects endorsed as old for choice versus fixed conditions within-subjects for each testing day. Finally, to determine if there was an influence of consolidation on choice-memory benefits, we compared forgetting rates across the immediate and 24-hour memory test (Leib Litman & Davachi, 2008). Forgetting was determined as a proportional difference in recognition memory across tests [(Day 1– Day 2)/(Day 1)] for each condition, separately. Forgetting rates were submitted to a paired t-test with condition (choice, fixed) as a within-subjects factor.

### Study 2: fMRI data acquisition and preprocessing

Functional imaging data were acquired on a Siemens Allegra 3T head-only scanner using echo planar imaging (TR=2000 ms; 34 contiguous slices; voxel size=3mm isometric). Slices were positioned parallel to the anterior commissure-posterior commissure and include coverage of our regions of interest (i.e., striatum, hippocampus, PRC). Our planned analyses focused on the pre and post-ratings task. Pre- and post-ratings functional MRI runs consisted of 308 volumes. We also collected a high-resolution T1-weighted anatomical scan (MPRAGE, voxel size = 1mm isotropic) for use in spatial normalization. Before fMRI preprocessing, data were inspected on custom software for head motion and scanning artifacts. Data were analyzed only if they exhibited <3.0mm motion (absolute maximum). Slice acquisitions with isolated transient noise artifacts were replaced with interpolated data from neighboring time points. fMRI preprocessing was then performed using FEAT (for FMRIB fMRI Expert Analysis Tool) version 6.00 as implemented in FSL version 5.0.2.1. The first four scans of each run were discarded for signal saturation. Images were skull-stripped, realigned, intensity normalized, and spatially smoothed with a 5.00 mm full-width half-maximum kernel, and subjected to a high pass filter (Gaussian-weighted least-squares straight line fitting set to 50.0s). fMRI images were transformed to standard space by first registering images to a subject-specific high-res anatomical image. We then applied a transformation matrix derived by a nonlinear transformation with a 10-mm warp resolution and 2 mm isotropic voxel resolution from the subject-specific anatomical image to an MNI standard-space image, as implemented in FMRIB Non-Linear Registration Tool.

### Study 2: Characterizing Pre- and Post-encoding hippocampal-PRC network coupling

To characterize post-encoding neural markers of consolidation, we measured functional coupling between hippocampus and perirhinal cortex during the ‘ratings task’ using a ‘background connectivity’ approach. This approach has been used in previous publications to measures low-frequency (state) changes in network interactions by removing task or trial evoked activation related to the ratings task (Al-Aidroos, Said, & Turk-Browne, 2012; K. Duncan, Tompary, & Davachi, 2014). Previous work in our laboratory has used this approach to assess how post-encoding changes in network coupling during an orthogonal (i.e. math) task related to subsequent memory (Tompary et al., 2015). We believe this approach is advantageous to a resting-state approach, as it better controls for explicit rehearsal of study materials. We first removed activity due to individual events in the pre- and post-encoding ratings task, separately. We modeled each event using a GLM that included separate regressors modelling hiragana characters that had appeared in (1) the choice trials, (2) the fixed trials, and (3) those that were not used in the memory encoding session, as well as their temporal derivatives. Each event was modeled with an event duration of 3 seconds convolved with a double-gamma hemodynamic response function. Data were also pre-whitened prior to analysis. We then extracted the time series from the residuals of these models from our regions of interest in the hippocampus and PRC (Figure 3, Left), separately for the right and left hemisphere. The hippocampus and PRC were defined using probabilistic atlases thresholded to 50% overlap. The hippocampus was defined from Harvard-Oxford Subcortical atlas (www.fmrib.ox.ac.uk/fsl/fslview). The PRC was defined from a probabilistic atlas generated by the Memory Modulation Lab (Ritchey, McCullough, Ranganath, & Yonelinas, 2017). Simple regressions were run in Matlab for ROI pairs of interest for each subject individually.

### Study 2: Analysis

To determine whether there were differences in functional coupling between the hippocampus and PRC after encoding, we compared r-scores from the pre-and post-encoding ratings task. Functional coupling from each ROI pair was compared using a paired t-test with the ratings session (Post, Pre) as a within-subjects factor. In our prior work, we found there was greater engagement of the striatum on choice versus fixed trials during encoding (Murty et al., 2015) To determine whether there were relationships between this choice-related striatal activity and post-encoding functional coupling, we ran correlations between striatal activation from the memory encoding phase and hippocampal-PRC coupling (Post > Pre). We only ran this analysis on pairs of hippocampal-PRC rois that showed significant differences in comparisons of post- and pre-encoding functional coupling. Striatal activations were extracted in response to the cues indicating choice and fixed trials during memory encoding. Details on how these values were estimated and extracted can be found in our previously published work (Murty et al., 2015). Correlations across conditions were tested using a fisher r-to-z transform. Finally, we ran a series of analysis to investigate how postencoding functional coupling was related to 24-hour memory. First, we tested whether differences in post-encoding functional coupling (Post>Pre) was related to 24-hour recognition memory for the choice condition using a simple regression. Then, to determine if our previously reported relationship between choice striatal activation and 24-hour memory was explained by differences in hippocampal-PRC functional coupling we ran the mediation package on R. Significance was tested using a non-parametric bootstrapping method, and we report on the average causal mediation effect.

## Results

### Study 1

#### Object Memory Performance

Participant’s memory was tested immediately and after a ~24-hour delay for the objects appearing in the choice and fixed conditions during encoding. Participants memory performance was significantly above chance both during the immediate memory test (Hits > FA; choice: t(32)=19.7, p<0.001; fixed: t(32): 18.9, p<0.001) and delayed memory test (Hits > FA; choice: t(32)=14.1, p<0.001; fixed: t(32): 10.5, p<0.001). Participants showed greater memory for objects in the choice versus fixed condition during the immediate memory test (Choice>Fixed; t(32) = 3.06, p=0.005) and delayed memory test (Choice>Fixed, t(32)= 4.38, p<0.001).

#### Forgetting rates

To test whether making decisions influenced memory consolidation, we examined forgetting rates by comparing memory performance across the immediate and 24-hour memory tests. Memory performance significantly declined across the 24-hour delay in the choice and fixed conditions (choice: t(32)=3.81, p<0.001; fixed: t(32)=9.69, p>0.001). However, forgetting rates were significantly lower for objects presented in the choice versus fixed condition (Figure 2; t(32)=2.36, p=0.02). Thus, chosen objects (or objects presented in the choice condition) showed better long-term memory retention, suggesting that processes occurring after encoding were stabilizing these representations.

**Figure 2.**
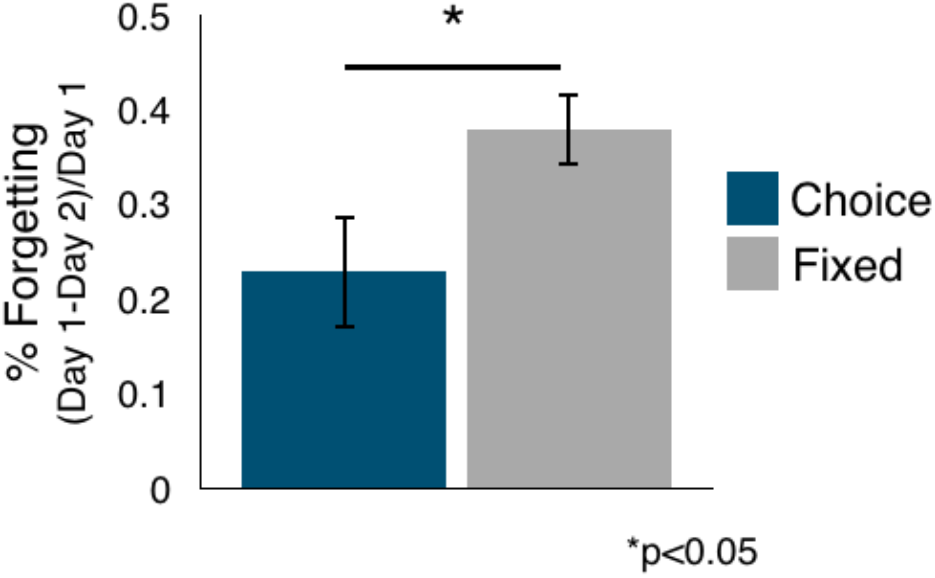
Active decision-making during encoding reduces forgetting. Participants showed decreased forgetting across the immediate and 24-hour memory test for objects encoded in the choice condition (blue) compared to the fixed condition (grey, p<0.05). Error bars represent standard error of the mean of each condition.

### Study 2

#### Changes in hippocampal-PRC coupling after encoding

We compared differences in functional coupling between the hippocampus and PRC before and after encoding, given that functional coupling between these regions has previously been associated with object-based memory consolidation (Vilberg & Davachi, 2013). Functional coupling was significantly greater following encoding between the right hippocampus and PRC (Figure 3; t(19)=3.43, p=0.003). This effect was trending towards significance between the left hippocampus and PRC (Figure 3; t(19)=1.79, p=0.09).

**Figure 3.**
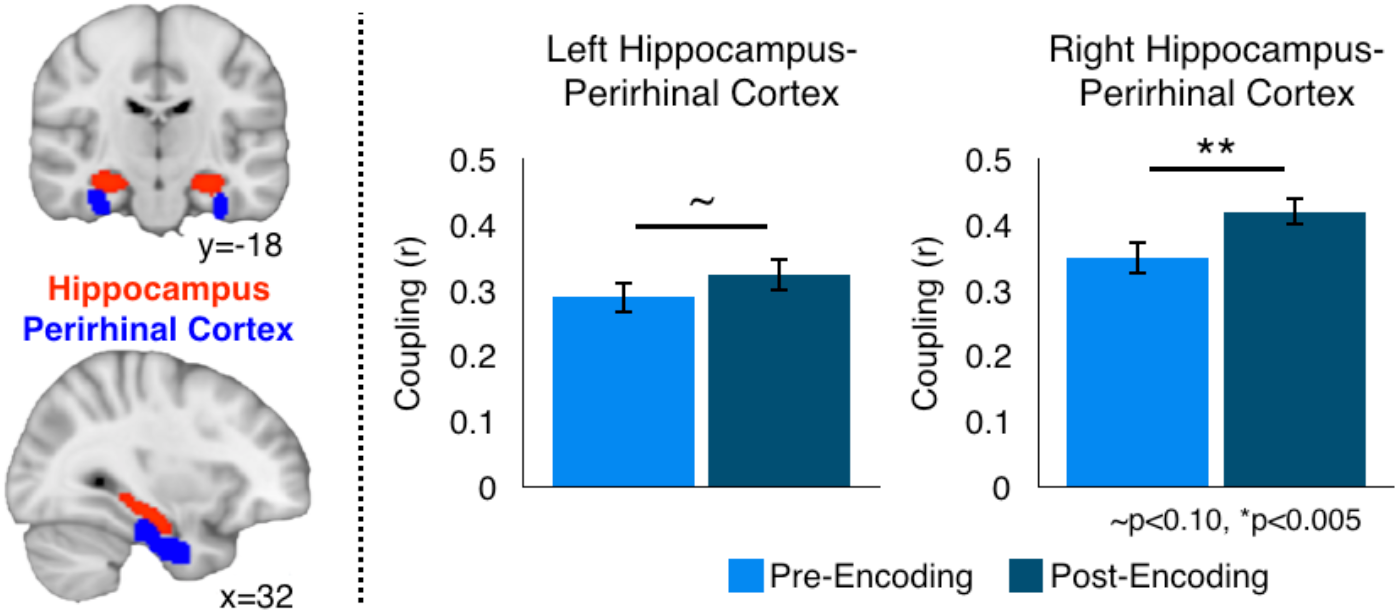
Encoding increases functional coupling between hippocampal-PRC. Regions of interest were defined in the hippocampus and perirhinal cortex (left). Functional coupling between the hippocampus and PRC was increased post-encoding in the right hemisphere (p<0.005), and a similar trend was seen in the left hemisphere (p<0.10). Error bars represent standard error of the mean.

#### Relationships between encoding-related striatal activity and post-encoding hippocampal-PRC interactions

We previously showed that choice versus fixed cues increased activation in the left striatum and was related to choice condition memory (Murty et al., 2015). Here, we tested whether striatal activation during encoding was related to post-encoding changes in functional coupling between right hippocampus and PRC. We found that striatal activation in response to choice cues was positively associated with increased post-encoding coupling of the right hippocampus and PRC (Figure 4; r(18)=0.51, p=0.02). There was no such relationship with striatal activation in response to fixed cues (Figure 4; r(18)=-0.12, p=0.60), and correlations were significantly greater for striatal activations in response to choice versus fixed cues (p=0.04, Z = 2.05).

**Figure 4.**
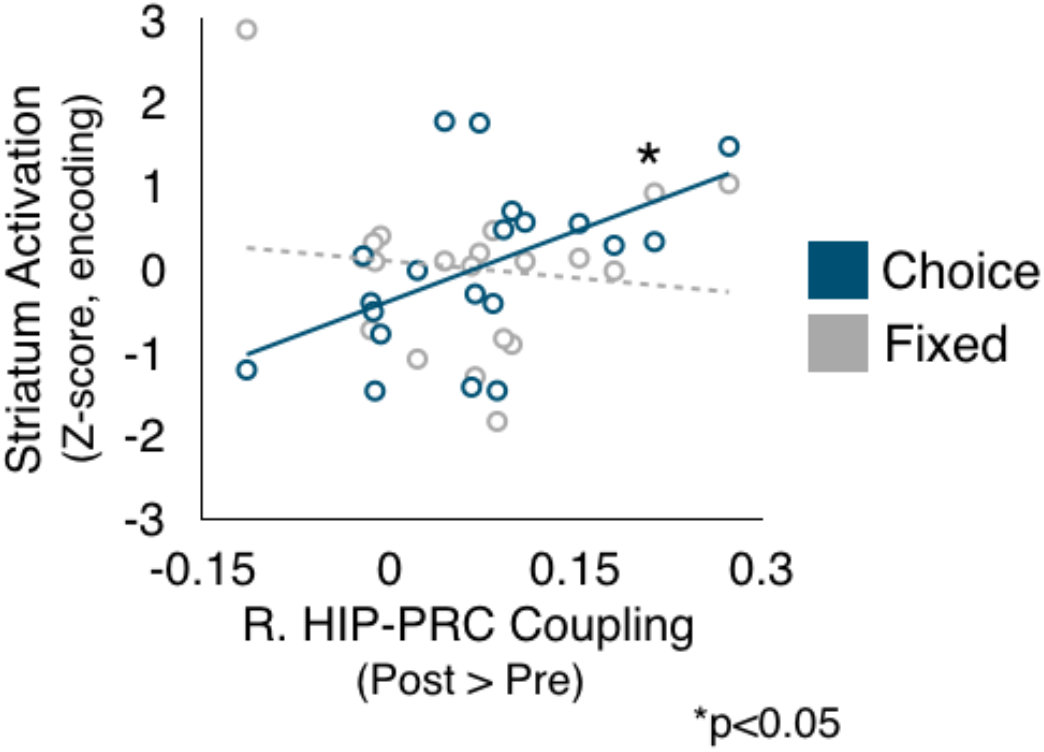
Choice-related striatal activation during encoding is associated with increased functional coupling between hippocampal-PRC after encoding. Striatal activation during the choice encoding condition (blue) was significantly related to increase right hippocampal-PRC coupling post encoding (p<0.05). No such relationship was seen for the fixed encoding condition (gray). Solid lines indicate significance

#### Relationships between 24-hour choice memory and post-encoding hippocampal-PRC interactions

We next tested whether post-encoding hippocampal-PRC interactions were related to memory for objects in the choice condition. Individuals that showed greater post-encoding increases in right hippocampal-PRC coupling had better 24-hour memory for objects encoded in the choice condition (r(18)=0.45, p<0.05). This relationship was not significant for objects encoded in the fixed condition (r(18)=0.37, p=0.11), but these correlations did not significantly differ from each other (p=0.55, Z=0.59).

Previously, we showed that choice-related striatal activation was also related to 24-hour choice memory (Murty et al., 2015). In a final analysis, we test the explanatory role of hippocampal-PRC post-encoding coupling in accounting for relationships between striatum and memory. We found that changes in hippocampal-PRC coupling significantly mediated the relationship between choice striatal activity 24-hour choice memory (p<0.05, one-tailed; Figure 5, Left). Critically, the model testing a mediating role for choice memory on the relationship between striatal activation and post-encoding connectivity was non-significant (p=0.17), and a model testing a mediating role for striatal activation on the relationship between post-encoding connectivity and memory was non-significant (p=0.42).

**Figure 5.**
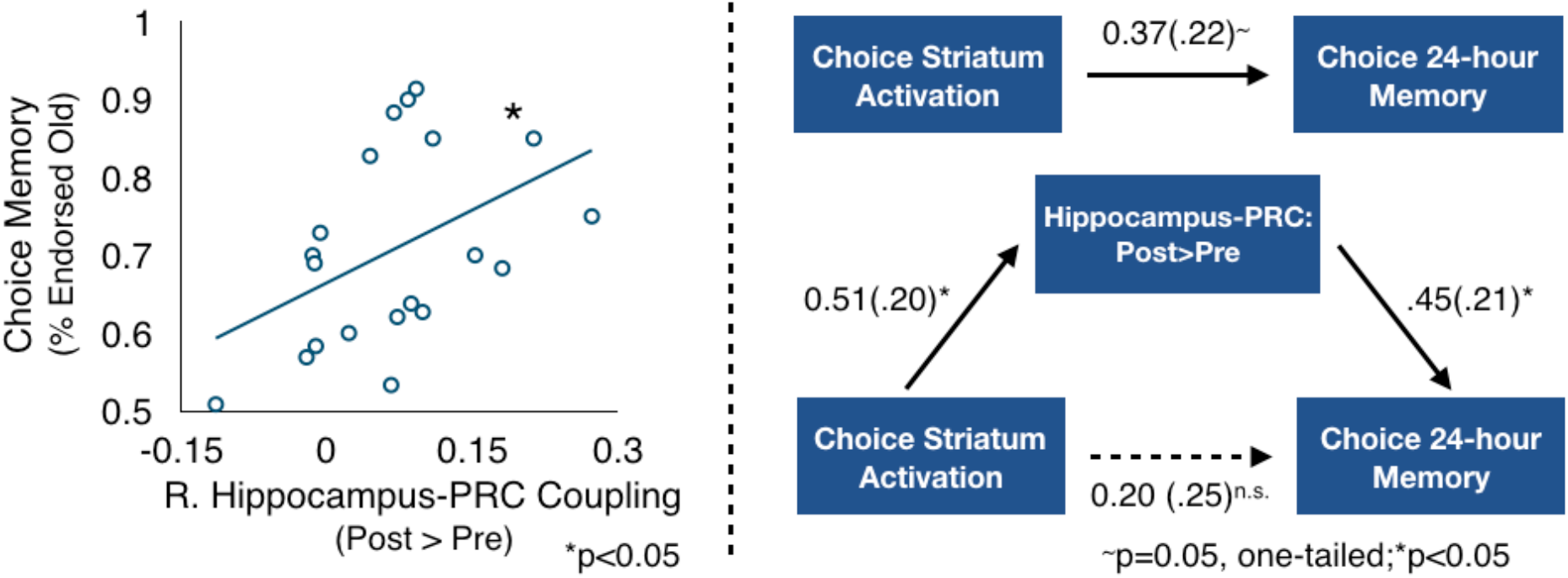
Changes in Hippocampal-PRC coupling is related to 24-hour memory for choice items. Changes in functional coupling between the hippocampus and PRC post versus pre-encoding significantly predicted 24-hour memory for objects in the choice condition (left, p<0.05). Changes in functional coupling between the hippocampus and PRC post versus pre-encoding mediated the relationship between choice striatal activation and 24-hour memory for objects in the choice condition (right, p<0.05, onetailed).

## Discussion

In the current study, we characterized how post-encoding mechanisms of consolidation influence interactions between decision-making and episodic memory. We found that both behavioral and neural measures of consolidation were related to decision-related enhancements in episodic memory. First, in a behavioral study, we showed that items participants actively decided to encode compared to items selected by the experimenter showed an enhanced resistance to forgetting over a 24-hour delay. Second, we found increased post-encoding interactions between hippocampus and PRC and these enhancements were related to both choice-related striatal activation during encoding and subsequent 24-hour memory. Together, these findings support a model in which decision-making enhances episodic memory, in part, by upregulating post-encoding consolidation processes.

A variety of different affective contexts have been shown to enhance memory consolidation. Rodent and human research, alike, has shown that environmental threat, novelty, and reward processing all increase memory consolidation (McGaugh, 2004; Miendlarzewska et al., 2016; Wang & Morris, 2010), resulting in a resistance to forgetting for objects encoded in these contexts. Here, we show that the simple act of making an arbitrary decision, in the absence of explicit incentives, resulted in a behavioral profile consistent with greater consolidation. In our task, many other features of encoding that influence episodic memory were matched across conditions, including viewing time, motor demands, and the content of memoranda (Murty et al., 2015). However, individuals still showed a resistance to forgetting of information that was ‘chosen’. Giving individuals the opportunity to make decisions has previously been associated with both increased valuation and engagement of mesolimbic dopamine systems (Coppin et al., 2014; Izuma et al., 2010; Izuma & Murayama, 2013; Leotti & Delgado, 2011, 2014; Sharot, Martino, & Dolan, 2009). These findings raise the interesting idea that decision-making may increase consolidation by generating an affective context, and inducing consolidation mechanisms like those induced by reward, novelty, and threat.

In line with this interpretation, we found relationships between striatal engagement—a key node in the mesolimbic network associated with valuation—and neural markers of post-encoding. Specifically, striatal engagement during active decision-making, but not when selection was determined by the experimenter selection, was related to enhanced post-encoding coupling of the hippocampus and PRC. Systems-level memory consolidation is thought to rely on post-encoding interactions in which memory traces stored in hippocampus are distributed to cortical regions associated with the sensory content of memoranda (McClelland et al., 1995; Nadel et al., 2000). The PRC, an anterior portion of medial temporal lobe cortex, is preferentially activated by object images (compared to other visual categories) and is important for object memory (Awipi & Davachi, 2008; Buckley, 2005; Davachi, 2006; Davachi et al., 2003; Graham et al., 2010; Liang & Preston, 2017; L. Litman et al., 2009; Ranganath, 2010; Staresina et al., 2011), such as those used as memoranda in this study. One prior paper also demonstrated that hippocampal-PRC connectivity is a biomarker for enhanced memory consolidation for object images (Vilberg & Davachi, 2013). Thus, we propose that striatal engagement during active choice bolsters information transfer between hippocampus and cortical regions. Further, we show that these processes relate to enhanced memory. Post-encoding increases in hippocampal-PRC interactions predicted memory in the choice condition, and accounted for our previous demonstrated relationships between mesolimbic engagement and episodic memory (Murty et al., 2015).

Our findings extend prior literatures demonstrating that post-encoding hippocampal-cortical interactions support memory consolidation (Schlichting & Preston, 2016; Tambini et al., 2010; Tompary et al., 2015), by showing that regions within the mesolimbic system may engage systems-level consolidation. A growing body of rodent research has demonstrated that dopamine activation supports memory consolidation by stabilizing synaptic plasticity within the hippocampus (Huang & Kandel, 1995; Li, Cullen, Anwyl, & Rowan, 2003b). Here, we show that mesolimbic activation may support memory consolidation not only by stabilizing processes within the hippocampus but also supporting post-encoding hippocampal interactions with cortical regions. This interpretation dovetails well with prior work from our laboratory associating reward-memory benefits with post-encoding interactions between the hippocampus and sensory cortex (Murty et al., 2017), as well as generalization of consolidation-related memory benefits across sensory categories (Patil et al., 2017). Critically, our current findings and previous work do not directly measure mesolimbic dopamine activation, but rather induce behavioral contexts that have previously been associated with dopamine activation in rodent and human studies. Future studies incorporating PET imaging and/or drug manipulations will be necessary to fully test this proposed mechanism.

Our findings provide a novel mechanism by which decision-making influences episodic memory via enhancing post-encoding consolidation for selected objects. While the neural mechanisms underlying episodic memory and decision-making have often been studied independently, recent efforts have begun to integrate knowledge across these fields (Bornstein, Khaw, Shohamy, & Daw, 2017; Bornstein & Norman, 2017; K. D. Duncan & Shohamy, 2016; Gershman & Daw, 2017; Murty et al., 2016; Shadlen & Shohamy, 2016). A large focus of this literature has been to investigate how episodic memories contribute to later adaptive decision-making. Here, we provide evidence for a reciprocal interaction in which decision making promotes subsequent episodic memory by facilitating systems-level consolidation to stabilize decision outcomes in memory. These processes represent a highly adaptive mechanism by which information that is actively acquired is given prioritization in long-term memory.

